# Individual variability in neural representations of mind-wandering

**DOI:** 10.1101/2024.01.20.576471

**Authors:** Aaron Kucyi, Nathan Anderson, Tiara Bounyarith, David Braun, Lotus Shareef-Trudeau, Isaac Treves, Rodrigo M. Braga, Po-Jang Hsieh, Shao-Min Hung

**Affiliations:** Department of Psychological & Brain Sciences, Drexel University, Philadelphia, PA – USA; Department of Neurology, Northwestern University, Chicago, IL – USA; Department of Brain and Cognitive Sciences and McGovern Institute for Brain Research, Massachusetts Institute of Technology, Cambridge, MA – USA; Department of Psychology, National Taiwan University, Taipei – Taiwan; Waseda Institute for Advanced Study, Waseda University, Tokyo – Japan

## Abstract

Mind-wandering is a frequent, daily mental activity, experienced in unique ways in each person. Yet neuroimaging evidence relating mind-wandering to brain activity, for example in the default mode network (DMN), has relied on population-rather than individual-based inferences due to limited within-individual sampling. Here, three densely-sampled individuals each reported hundreds of mind-wandering episodes while undergoing multi-session functional magnetic resonance imaging. We found reliable associations between mind-wandering and DMN activation when estimating brain networks within individuals using precision functional mapping. However, the timing of spontaneous DMN activity relative to subjective reports, and the networks beyond DMN that were activated and deactivated during mind-wandering, were distinct across individuals. Connectome-based predictive modeling further revealed idiosyncratic, whole-brain functional connectivity patterns that consistently predicted mind-wandering within individuals but did not fully generalize across individuals. Predictive models of mind-wandering and attention that were derived from larger-scale neuroimaging datasets largely failed when applied to densely-sampled individuals, further highlighting the need for personalized models. Our work offers novel evidence for both conserved and variable neural representations of self-reported mind-wandering in different individuals. The previously-unrecognized inter-individual variations reported here underscore the broader scientific value and potential clinical utility of idiographic approaches to brain-experience associations.

## Introduction

The brain’s patterns of spontaneous activity are dynamic, yet highly organized, even during task-free periods when external inputs are unchanging.^1^ Mind-wandering, often defined as unconstrained, self-generated thoughts that are independent of stimuli and tasks,^2^ is a prevalent form of mental activity during these task-free periods and in daily life in general.^3,4^ What people naturally think about, and how thoughts dynamically unfold, are fundamental to cognitive function and mental health.^5^ Recent studies have begun to delineate how mind-wandering arises from spontaneous brain activity and have emphasized a key role of the default mode network (DMN).^5–8^

The most commonly employed method to study mind-wandering is experience-sampling, wherein people respond to “thought probes” about their subjective level of focus on a cognitive task^9^ (thereby defining mind-wandering as off-task thought^6^). Experience-sampling has been combined online with neuroimaging, revealing an association between mind-wandering with DMN activation^10–14^ and broader patterns of whole-brain functional connectivity.^14–16^ These neuroimaging experiments have typically included ∼30-40 total thought probes in a single 1-2 hour session.^10–15,17–21^ Such limited within-subject sampling has led studies to almost exclusively rely on population-level inferences, which harbor the implicit assumption that mappings between brain activity and self-reported mind-wandering are conserved across individuals.

Multiple theoretical and empirical considerations challenge this assumption. First, the content and form of mind-wandering varies substantially across individuals,^22,23^ and different subtypes of ongoing thought have distinct neural bases.^17,24^ The specific experiences that tend to occur when different people report off-task thought likely contribute to variable brain-experience mappings. Second, focal lesions within DMN-related regions^25,26^ primarily impact the content of thought without abolishing the ability to engage in mind-wandering.^8^ This points to the possibility of degenerate neural mechanisms,^27^ or multiple sets of activity patterns that can give rise to similar self-report outcomes. Third, recent neuroimaging evidence highlights individual variability in neural representations of subjective experiences,^28^ especially when experiences are self-related.^29^ Taken together, these considerations motivate an idiographic (personalized) approach that prioritizes rich, within-person sampling to enable individual-level inferences and tests of group-to-individual generalizability.^30^

Importantly, even if there are conserved neural patterns linked to mind-wandering across individuals, neuroimaging studies have not been optimally conducted to detect such patterns. Following common practices in the field, studies have relied on inter-subject registration of individual brain images to common-space templates, a method that inevitably results in cross-subject misalignment.^31^ Recent studies using “precision functional mapping”^32,33^ have established that the detailed functional anatomy of the DMN and other networks conforms to the unique cortical folding patterns of different brains.^34–36^ Estimating networks within individuals, which can be achieved reliably with dense sampling,^33^ may unravel why associations between DMN activation and mind-wandering have not always been replicated^15^ or found consistently within all individuals.^13^

Here we sought to determine whether and how neural patterns linked to mind-wandering may be distinct across individuals. We leveraged data from individuals who underwent multiple sessions of functional magnetic resonance imaging (fMRI) where experience-sampling was embedded into a “resting state” (simple visual fixation) condition, resulting in over 300 thought probes per subject. This paradigm allowed us to examine, with rich individual-level detail, the relationship between natural fluctuations in inner experience and spontaneous brain activity. We applied precision functional mapping and hypothesized that mind-wandering reports would relate (a) reliably to activation of the DMN, including temporal alignment to subjective reports; and (b) unreliably to activation of networks implicated in specific thought features that tend to vary across individuals. We further applied idiographic predictive modeling based on functional connectivity to explore individual-level specificity of whole-brain representations of mind-wandering. We formally compared predictive neural patterns between subjects and tested group-to-individual generalizability using recently developed population-level neuroimaging models of constructs related to mind-wandering.^16,37^

## Results

### Mind-wandering fluctuations in multi-session fMRI

Three subjects (S1-3) each completed 6 fMRI sessions across different days. In one session, they performed functional localizer task runs that we used for precision functional mapping. In each of the other 5 sessions, they performed 7-11 runs (∼8 mins/run) of an experience-sampling task that we used for main analyses. The experience-sampling condition involved a simple visual fixation task, coupled with intermittent thought probes appearing every 45-90 seconds (**Fig. 1a**). During thought probes, subjects indicated the degree to which they were focused on the fixation task on a 1-8 Likert scale (mind-wandering rating). Alternatively, subjects could respond that they were distracted but were not mind-wandering (e.g. attending to an external stimulus); for the purposes of the present study, we discarded these distraction trials. If subjects reported that they were not focused on the task (rating of 5 or higher on 1-8 scale), then the trial was deemed “mind-wandering,” and they responded to a series of additional items about thought content and form (see **Methods**).

**Figure 1.**
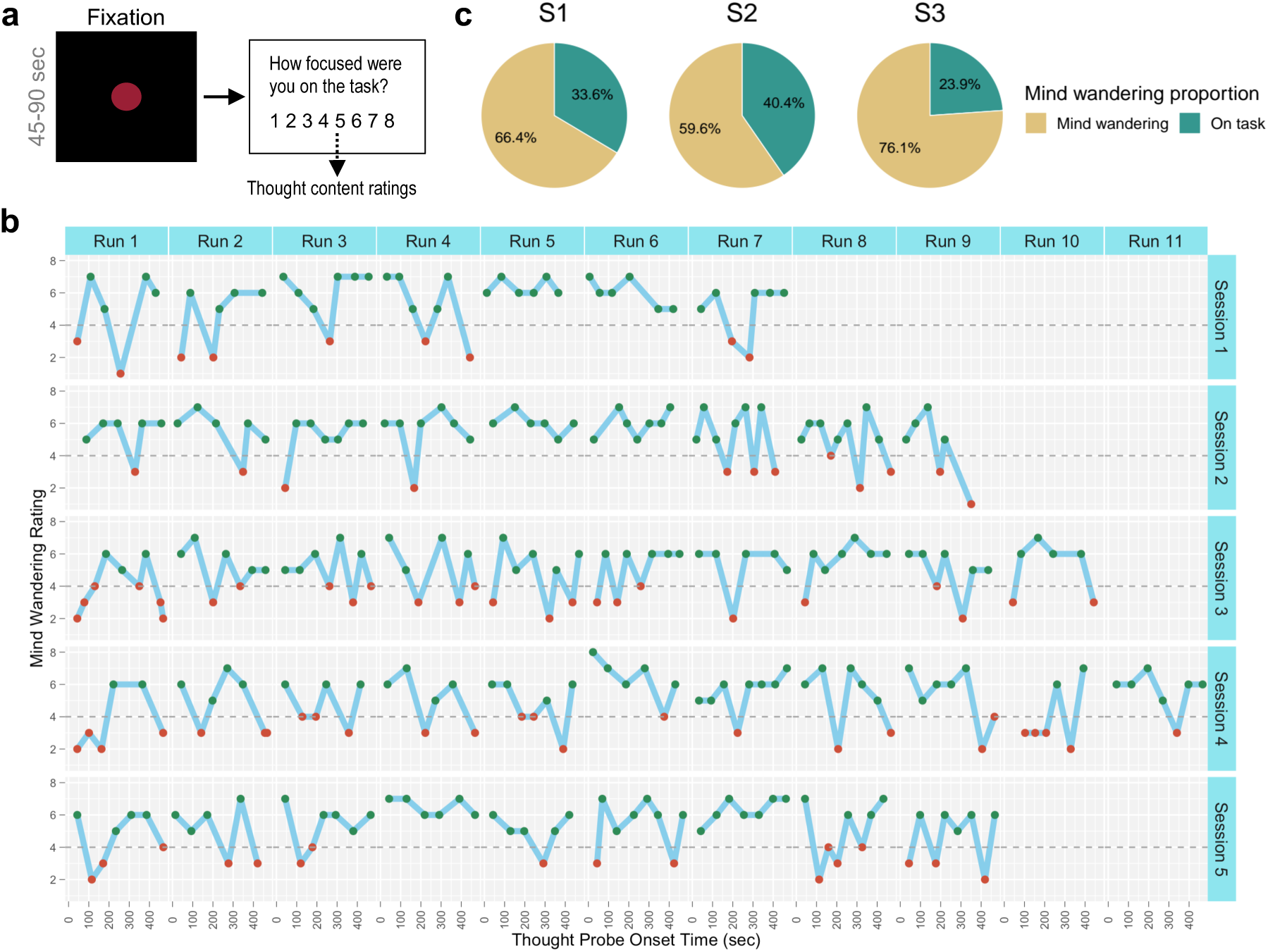
Experience-sampling paradigm and mind-wandering ratings. **a)** In fMRI runs involving experience-sampling, subjects were instructed to stare at a red fixation circle. They were intermittently probed every 45-90 seconds with the question (Q1): “How focused were you on the task?”, accompanied by an 8-point Likert scale. A focus rating of 5 or greater was deemed a “mind-wandering trial” and triggered an additional set of questions asking about thought contents. During data analyses, ratings were reverse-coded so that a higher rating indicated more mind-wandering. **b)** Example time series of single-subject mind-wandering ratings. The plots show subject S1’s trial-by-trial responses to Q1 across runs within each MRI session. Green and red markers, respectively, indicate trials that were labeled as mind-wandering and non-mind-wandering. Trials in which a different response than 1-8 was submitted (see Methods) are not shown. **c)** Proportion of trials in which mind-wandering was reported in each subject. **d)** Proportions of different thought categories reported during mind-wandering in each subject (see also Fig S1a).

Each subject reported a wide range of task-focus ratings within and across runs and sessions (example shown in **Fig. 1b**). After discarding distraction trials, 256 or more trials per subject were retained (S1: n=318; S2: n=256; S3: n=316), and over 50% of these trials were consistently considered “mind-wandering” based on rating binarization (S1: 66.4%; S2: 59.6%; S3: 76.1%) (**Fig. 1c**). Depending on the subject, mind-wandering ratings were not correlated, or were weakly correlated, with head motion, time within runs, and trial order within sessions (**Table S1a**). This suggests that there were no consistent associations across subjects between mind-wandering reports and nonspecific factors related to drift over time, such as arousal. However, we present relevant control analyses within subjects where appropriate below.

As expected, when subjects reported mind-wandering, specific thought content and form varied across subjects. Predominant thought categories included visual and auditory imagery. Across trials, subjects reported a wide variety of specific mental activities (e.g. recalling, planning and imagining), perspectives (e.g. first-person vs. third-person), and emotions (see **Fig. S1a** for detailed breakdown of thought subcategories). Inter-subject comparison revealed that the dissimilarity between each pair of subjects was dependent on thought subcategory; for example, while S1 and S2 were similar in terms of content modality (**Fig. 1d**), they were dissimilar in terms of type of mental activity (e.g. planning vs. recalling) (**Fig. S1b**). Taken together, while mind-wandering was consistently reported frequently across fMRI sessions, specific content and form of thought varied across subjects.

### Activation of the DMN, estimated within individuals, is associated with mind-wandering

We next conducted within-individual analyses of the relationship between trial-by-trial variations in mind-wandering ratings and blood oxygen level dependent (BOLD) signal activation within the DMN. To precisely map the DMN within individuals, we used multi-session hierarchical Bayesian modeling (MS-HBM), a method for reliable parcellation of the cerebral cortex into a discrete set of labeled systems.^38^ We applied MS-HBM based on functional connectivity analysis of fMRI functional localizer data that was independent from the experience-sampling data (see **Methods**). Regardless of how the parcellation level was specified (**Fig. S2**), two DMN-related networks were consistently identified within each subject. These networks demonstrated inter-individual variability in spatial arrangements but had organizational motifs that were consistent with previously described fractionation of the DMN into default network A (DNa) and B (DNb)^34^ (**Fig. 2a**). The two networks neighbored one another and included prominent subregions within medial prefrontal, posteromedial, inferior parietal and lateral temporal cortices, while a distinguishing feature of DNa was that it was coupled to regions in parahippocampal cortex.^39^

**Figure 2.**
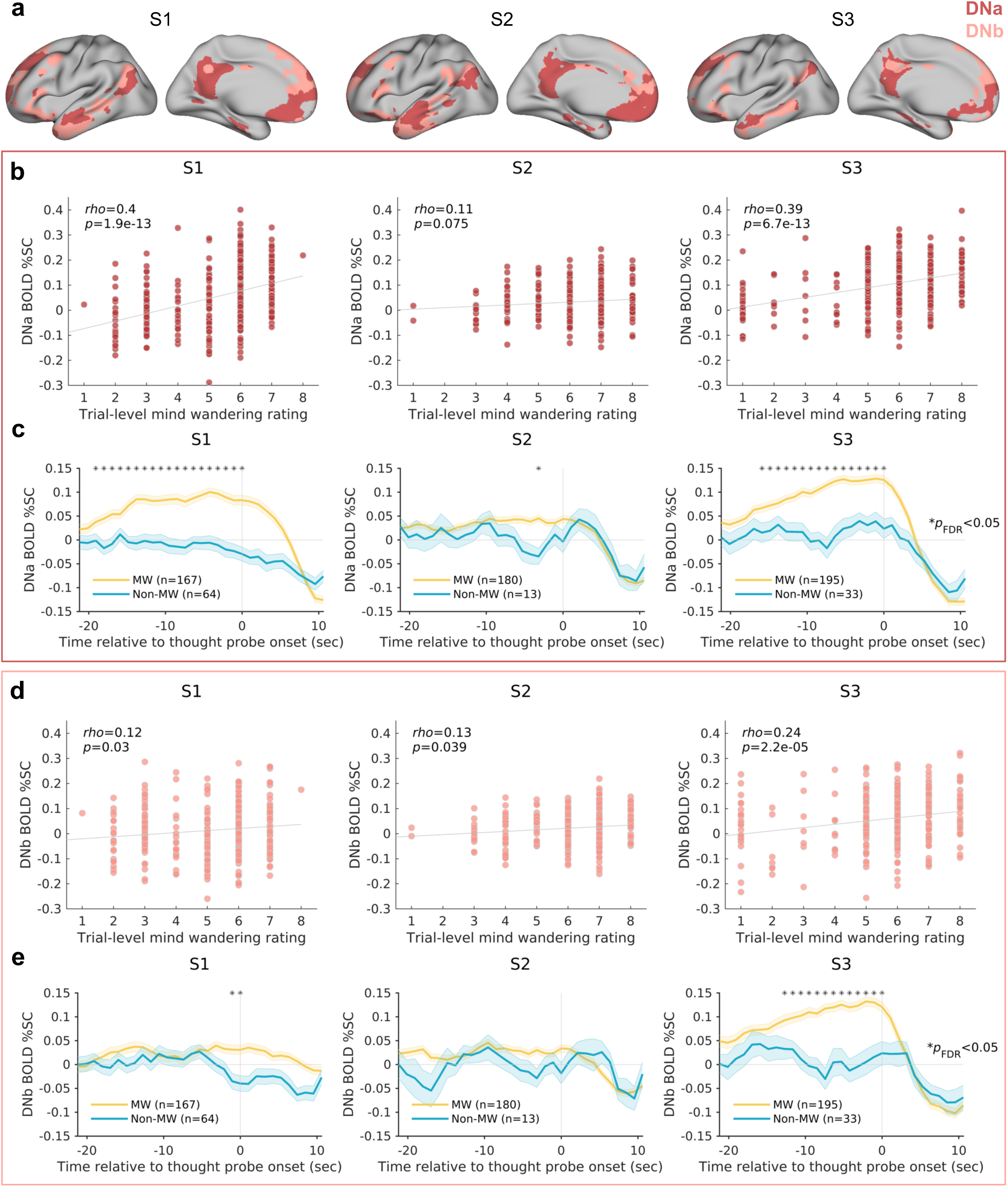
Activation of the DMN, estimated within individuals, is associated with mind-wandering. a) Two individually-estimated networks of the DMN (DNa and DNb) identified among 17 total cortical networks based on multi-session hierarchical Bayesian modeling applied to functional localizer fMRI data. b) Correlations between trial-by-trial mind-wandering ratings and median DNa blood oxygenation level dependent (BOLD) percent signal change (%SC) within 10-second pre-thought probe periods. **c)** Mean time series plots showing median BOLD %SC in DNa for mind-wandering trials (rating of 6 or higher) and non-mind-wandering trials (rating of 3 or lower) before and after thought probes. Asterisks indicate time points where there was a significant difference in BOLD %SC between trial types (Wilcoxon rank sum test; p<0.05, false discovery rate-corrected for number of samples within 20-second window prior to thought probes). Shaded error bars indicate standard error of the mean. **d)** Same as b) but for DNb. **e)** Same as c) but for DNb.

We extracted the median BOLD percent signal change (%SC) from DNa and DNb within the 10-sec periods prior to each thought probe. Within each of the three subjects, we found positive correlations between mind-wandering rating and pre-probe BOLD %SC in DNa, and these correlations were significant within two subjects (S1: Spearman’s *ρ* = 0.40, *p* = 1.9x10^-13^; S2: Spearman’s *ρ* = 0.11, *p* = 0.075; S3: Spearman’s *ρ* = 0.39, *p* = 6.7x10^-13^) (**Fig. 2b**; see **Table S1b** for control analyses). For DNb, significant positive correlations with mind-wandering were found within all three subjects (S1: Spearman’s *ρ* = 0.12; *p* = 0.03; S2: Spearman’s *ρ* = 0.13; *p* = 0.039; S3: Spearman’s *ρ* = 0.24; *p* = 2.2x10^-5^) (**Fig. 2d**; see **Table S1c** for control analyses), even though these correlations were weaker than the DNa correlations in two subjects (S1 and S3). For those two subjects, but not for S2, correlations between DNa and mind-wandering were significant over-and-above the contributions of DNb, as revealed by partial correlations between DNa BOLD %SC and mind-wandering while controlling for DNb BOLD %SC (S1: *ρ_partial_* = 0.44, *p* =3.3x10^-16^; S2: *ρ_partial_* = 0.039, *p* = 0.54; S3: *ρ_partial_* = 0.25, *p* = 6.0x10^-6^).

We repeated the correlation analyses using standard-space DMN subnetworks from a commonly used population-derived atlas (Yeo-Krienen 17-network parcellation).^40^ A standard-space subnetwork commonly referred to as the medial temporal lobe subsystem,^41^ which overlapped with DNa, showed positive correlations with mind-wandering in all subjects (**Fig. S3**). However, the MTL subsystem correlations were weaker than the within-subject DNa correlations, highlighting how precision functional mapping enhanced the ability to detect brain-experience relationships.

Importantly, the analyses presented so far focused on BOLD activation within 10-sec pre-probe periods, which is in accordance with prior work.^10,11,13^ However, the timing of spontaneous neural activations can vary relative to subjective reports of recalled experiences at the time of a thought probe.^42^ We therefore performed complementary analyses to explore the temporal dynamics of DNa and DNb activations relative to thought probe onsets. We computed BOLD %SC within each subnetwork for trials categorized as ‘pure’ mind-wandering (rating between 6 and 8) or non-mind-wandering (rating between 1 and 3), removing trials with intermediate ratings that may be selected when participants are less certain about their mental experience. This analysis revealed that within DNa, all three subjects displayed specific pre-probe timings where BOLD %SC was significantly greater for mind-wandering than non-mind-wandering trials (all *p_FDR_* < 0.05; Wilcoxon rank sum test) (**Fig. 2c**). In S1 and S3, mind-wandering-related DNa activation was found as early as 18-19 seconds preprobe and was sustained until thought probe onset. In S2, there was significant activation only at a time point ∼3 seconds preprobe. In all three subjects, DMN deactivation peaked ∼4-6 seconds after thought probe onset, a characteristic latency that was expected based on the typical timing of neurophysiological responses in the DMN^43^ and the hemodynamic delay of BOLD activity.^44^ Within DNb, only S1 and S3 (but not S2) exhibited preprobe timings where BOLD %SC was significantly greater for mind-wandering than non-mind-wandering trials (both *p_FDR_* < 0.05) (**Fig. 2c**).

In summary, these findings show that: (a) Within the 10-sec preprobe window, all subjects demonstrated positive relationships between DMN activation and mind-wandering, with two out of three subjects showing a preferential relationship with DNa relative to DNb; and (b) Spontaneous DNa activation was consistently related to mind-wandering within all subjects when accounting for variable timing of activation relative to subjective reports.

### Individual variability in functional networks activated and deactivated during mind-wandering

We have so far focused on region-of-interest analyses of DMN subnetworks, but mind-wandering has often been associated with activation and deactivation of other networks.^6^ We therefore next performed within-subject, whole-brain general linear model (GLM) analyses to search for additional brain regions associated with trial-by-trial mind-wandering ratings.

Unthresholded whole-brain statistical maps showing positive and negative associations with mind-wandering are shown in **Fig. 3a**, with suprathreshold significant regions outlined in black (family-wise error rate-corrected cluster-determining threshold: *Z* > 3.1; cluster-based *p* < 0.05). This conservative analysis (i.e., due to multiple comparisons correction across whole-brain voxels) revealed both significant positive and negative associations with mind-wandering in S1 and S3 but not in S2. In both S1 and S3, significant cortical regions were both within and outside of the DMN. In addition, there were significant associations between mind-wandering and activation within multiple subcortical and cerebellar regions (**Fig. 3b**). For example, in S1, left hippocampus and right amygdala, respectively, were positively and negatively associated with mind-wandering. In S3, right amygdala and bilateral thalamus, respectively, were positively and negatively associated with mind-wandering.

**Figure 3.**
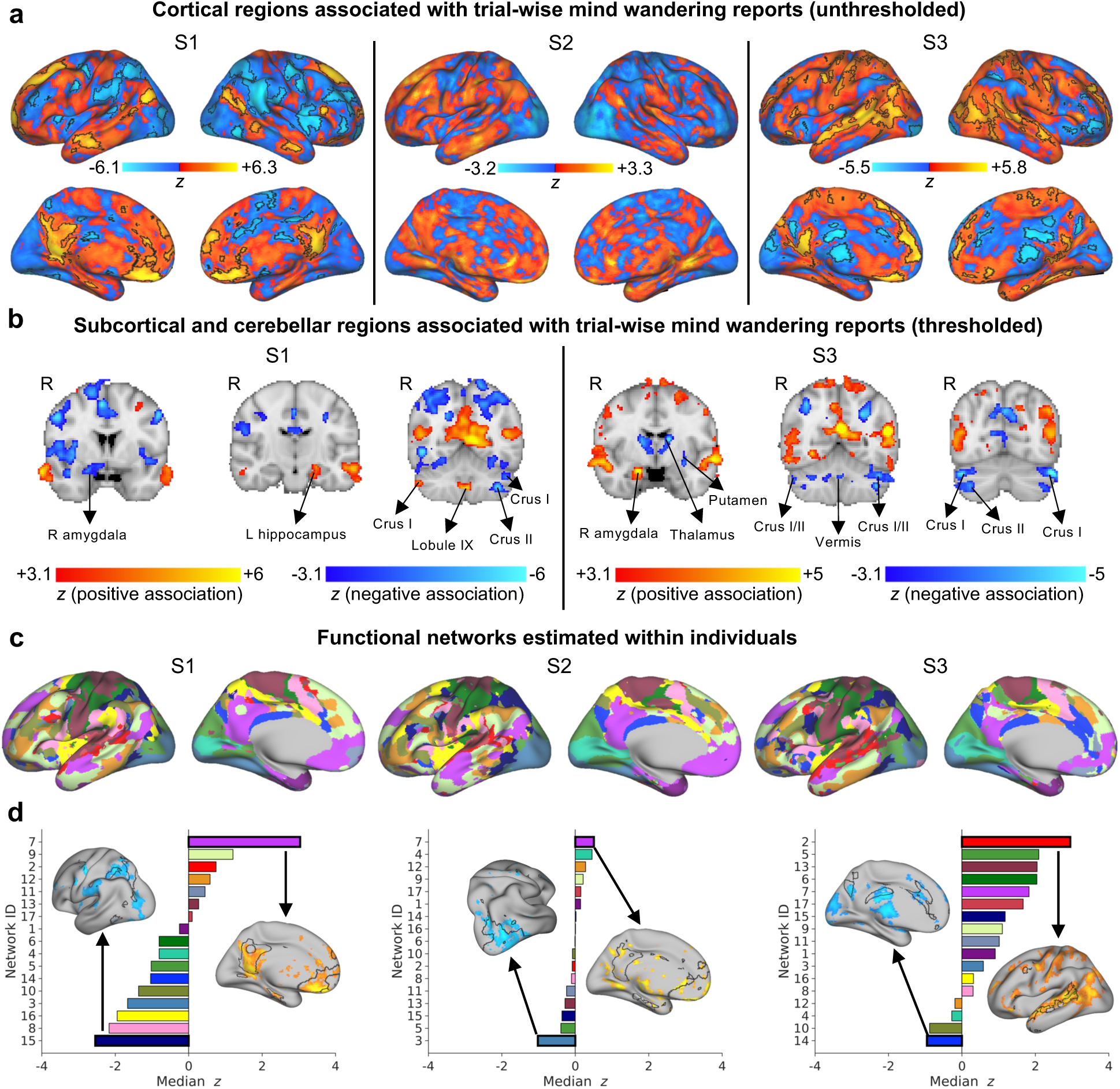
Individual variability in networks activated and deactivated during mind-wandering. **a)** Cortical surface plots of unthresholded statistical parametric maps showing regions positively (red-yellow) and negatively (blue-light blue) associated with mind-wandering during 10-second pre-thought probe windows. Maps are based on a whole-brain, within-subject general linear model (GLM) analyses combining all fMRI experience-sampling runs and sessions. Whole-brain corrected, statistically-significant clusters are outlined in black (FWE-corrected cluster-determining threshold: Z>3.1, cluster-based p<0.05). **b)** Representative coronal volumes in two subjects who showed significant clusters in subcortical and cerebellar regions. Statistical maps are thresholded to show only significant regions (FWE-corrected cluster-determining threshold: Z>3.1, cluster-based p<0.05). **c)** 17 personalized cortical networks identified with multi-session hierarchical Bayesian modeling applied to functional localizer fMRI data. **d)** Rank-ordered median z score (obtained from GLMs) within each one of the networks shown in b). Cortical surface plots are shown for the networks most positively and negatively associated with mind-wandering, respectively; statistical maps are individually thresholded to retain top voxels showing associations, and network outlines are shown in black.

To further characterize cortical areas with respect to networks estimated within individuals, we computed the median *z* statistic scores obtained from GLMs within 17 networks independently mapped with MS-HBM (**Fig. 3c**). We rank-ordered these networks based on strength of association with mind-wandering (**Fig. 3d**). For S1, the top (most positively associated) network was DNa, and the bottom (most negatively associated) network was consistent with the dorsal attention network (e.g. superior parietal lobule, frontal eye fields), a network that classically shows activity that is negatively correlated with the DMN.^45^ For S2, the top and bottom networks, respectively, were DNa and a network involving lateral visual regions. For S3, the top and bottom networks, respectively, were a network that included language-related regions (e.g. left lateral temporal cortex) and a network that was consistent with the salience network^46^ (e.g. ventral anterior insula, anterior mid- and posterior-cingulate cortex, and precuneus); the positive associations between mind-wandering and the language network were even stronger than associations with DNa or DNb.

In summary, extending analyses to other networks, there was substantial individual variability in brain activations and deactivations during mind-wandering. To further explore individual variability in broader neural representations at the level of functional network coupling, we next performed a data-driven, idiographic predictive modeling analysis.

### Idiographic connectome-based predictive modeling of mind-wandering

Prior work showed that dynamic BOLD functional connectivity can contribute variance to prediction of mind-wandering over-and-above BOLD activation.^12,14,15^ We recently applied connectome-based predictive modeling (CPM) to develop a whole-brain, multivariate, functional connectivity model of mind-wandering (defined as stimulus-independent and task-unrelated thought) where model training was performed using combined trials across subjects.^16^ Here, in our densely-sampled subjects, we performed functional connectivity analyses to address three major questions: (a) Can we generate successful predictions of mind-wandering from CPM performed fully within single subjects (i.e., idiographic CPM) (**Fig. 4**)? (b) If so, are predictive features similar or different across subjects (**Fig. 5)**? (c) Can mind-wandering be predicted within densely-sampled subjects using models generated from other densely-sampled subjects or from populations of subjects (**Fig. 6**)?

**Figure 4.**
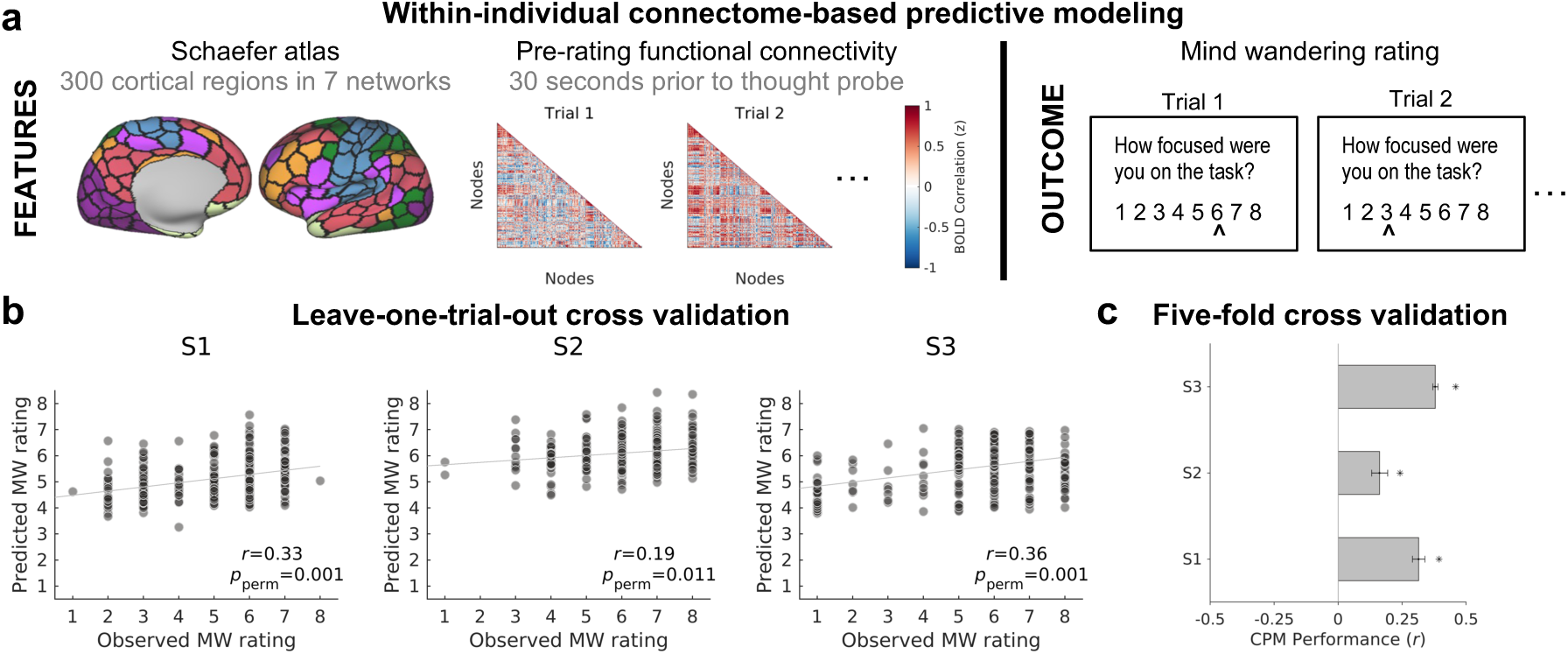
Within-individual connectome based predictive modeling (CPM) of mind-wandering. **a)** Features (left) for CPM included functional connectivity (correlated activity) between regions in a 300-region cortical atlas within 30-second windows prior to thought probes. The CPM outcome to be predicted (right) was mind-wandering rating. **b)** Correlation between predicted and observed mind-wandering within each subject based on CPM with leave-one-trial-out cross validation. **c)** Mean correlation between predicted and observed mind-wandering, based on CPM with five-fold cross validation (see also Table S1d). Error bars show standard deviation of correlations across 120 iterations of cross-validation.

**Figure 5.**
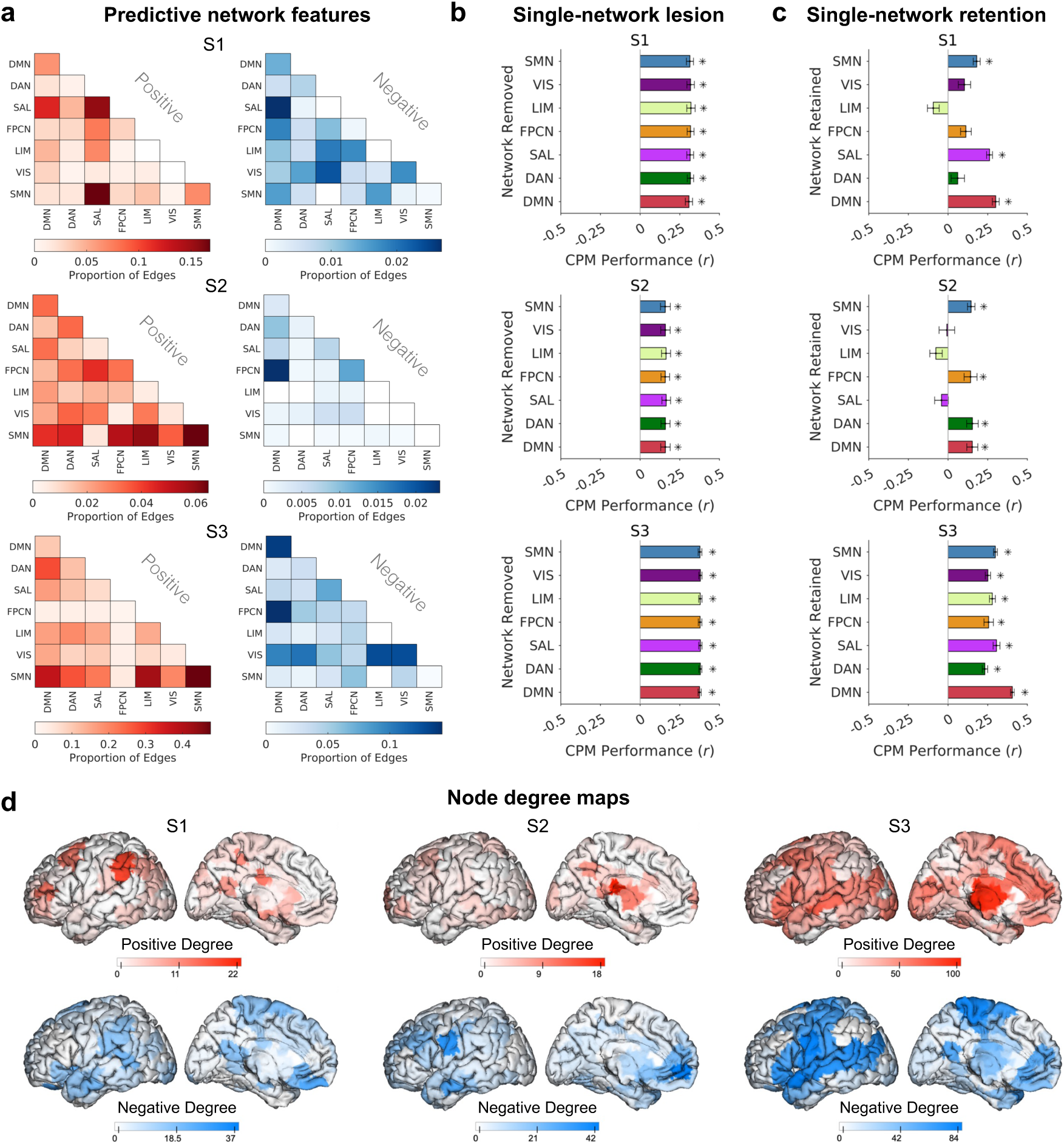
Individual variability in functional anatomy of features important for connectome-based predictive modeling of mind-wandering. **a)** Summaries of predictive network features that were selected in CPM to model and predict mind-wandering within each subject. For both positive and negative predictive region-pairs (edges), the proportion of total edges for each pair of networks in the Yeo-Krienen 7-network atlas is shown. **b)** Mean correlation between predicted and observed mind-wandering, based on CPMs with different networks virtually lesioned (i.e., all edges involving the ‘lesioned’ network removed for CPM training and testing). **c)** Mean correlation between predicted and observed mind-wandering, based on CPMs with all networks removed except for one (i.e., only edges involving the labelled network retained for CPM training and testing). Results shown in b) and c) are based on five-fold cross validation. **d)** Left hemisphere cortical maps showing node degree (i.e., number of total edge contributions) for regions that positively (red) and negatively (blue) contributed to CPMs within each subject. All error bars indicate standard deviation of correlations across 120 iterations of five-fold cross validation. DAN = dorsal attention network; DMN = default mode network; FPCN = frontoparietal control network; LIM = limbic network; SAL = salience network; SMN = sensorimotor network; VIS = visual network

**Figure 6.**
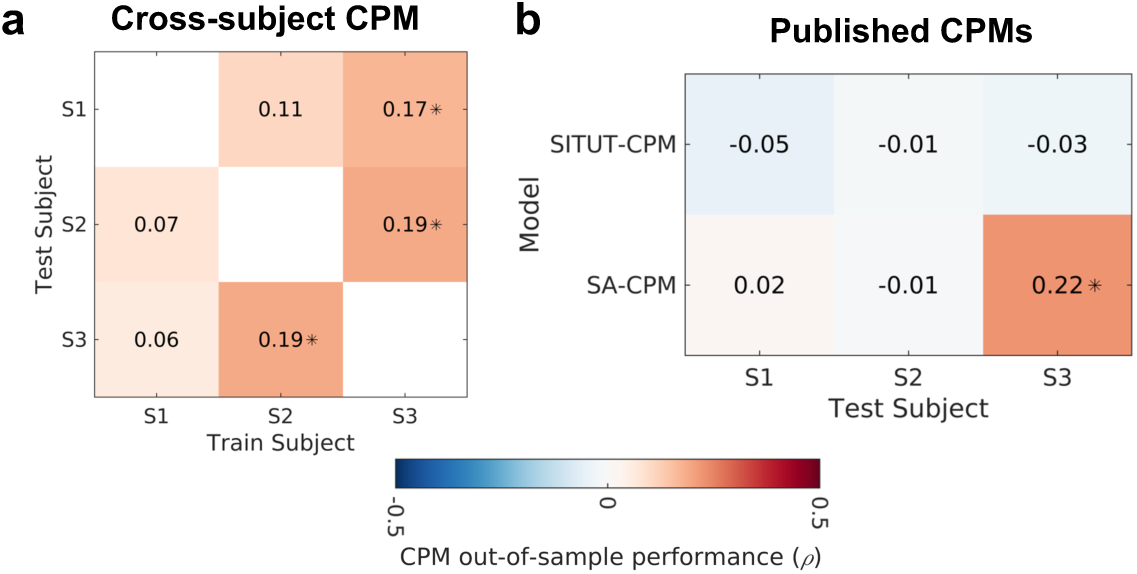
Predicting trial-wise mind-wandering from cross-subject and population-derived connectome-based predictive models. **a)** Spearman rank correlation coefficients for predicted versus observed mind-wandering using training data from subjects shown on x axis and test data from subjects shown on y axis. **b)** Spearman rank correlation coefficients for predicted versus observed mind-wandering using two previously published, population-derived CPMs applied to S1, S2 and S3 as test subjects. Asterisks indicate p<0.05. SITUT-CPM = stimulus-independent, task-unrelated thought CPM; SA-CPM = sustained attention CPM.

We performed idiographic CPM following procedures that closely matched our prior work.^16^ Briefly, we extracted functional connectivity (activity correlation) between all region-pairs in a 300-region cortical atlas within 30-second windows prior to thought probes (**Fig. 4a**). Within training data (a subset of trials), features were selected based on strength of positive or negative association with mind-wandering rating. Within independent, held-out (testing) trials, we compared predicted versus observed mind-wandering and used permutation tests to assess significance (see **Methods**).

Idiographic CPMs yielded significant prediction of mind-wandering within each subject when using cross-validation schemes of either leave-one-trial-out (S1: *r* = 0.33, *p_perm_* = 0.001; S2: *r* = 0.19, *p_perm_ =* 0.011; S3: *r* = 0.36, *p_perm_ =* 0.001) (**Fig. 4b**) or five-fold (S1: *r_mean_ =* 0.31*, p_perm_* = 0.001; S2: *r_mean_ =* 0.16*, p_perm_* = 0.025; S3: *r_mean_ =* 0.38*, p_perm_* = 0.001) (**Fig. 4c**) (see **Table S1d** for control analyses). The results were consistent when using the alternative Shen 268-region whole-brain atlas^47^ (**Table S2a**) and when using brain basis set modeling^48^ instead of CPM (**Table S2b**). These findings confirm that whole-brain, multivariate patterns of dynamic functional connectivity were predictive of within-subject mind-wandering, even notably in S2 who had not shown significant whole-brain level activation effects in the univariate GLM analysis (**Fig. 3a**). Having confirmed the success of idiographic CPM, we next compared the features that drove predictions across subjects.

### Idiosyncratic network features predict mind-wandering in different individuals

In each subject, a large number of positive edges (S1: n=940; S2: n=322; S3: n=4954) and negative edges (S1: n=3568; S2: n=2650; S3: n=14748) contributed to CPMs. To characterize these predictive features, we labelled each node in a region-pair (edge) that positively or negatively contributed to a CPM as belonging to 1 of 7 intrinsic networks in the population-based Yeo-Krienen^40^ cortical atlas. Each subject showed an idiosyncratic, complex pattern of predictive network features comprised of within- and between-network edges involving all 7 networks (**Fig. 5a**). For example, while DMN within-network and inter-network edges were included in both positive and negative features in all subjects, the specific edges involved were variable across subjects.

To further determine the contributions of each network to CPM, we performed computational lesion analyses, where all edges from a selected network were removed during feature selection. When single networks were lesioned, CPM performance always remained significant, regardless of network removed (all *p_perm_* < 0.05) (**Fig. 5b**), with minimal effects on performance compared to full models that included all networks. However, the impact of retaining only one network (i.e., computational lesioning 6 out of 7 networks) was more variable across subjects, with CPM performance remaining significant only for certain single-network models. In S1, S2, and S3, respectively, three (sensorimotor, salience, and DMN), four (sensorimotor, frontoparietal control, dorsal attention, and DMN), and all seven (sensorimotor, visual, limbic, frontoparietal control, salience, dorsal attention, and DMN) single-network models yielded significant performance (all *p_perm_* < 0.05) (**Fig. 5c**). This suggests that functional connectivity within some single networks alone is consistently sufficient for successful idiographic CPM performance (sensorimotor network and DMN), while the contributions from other networks are subject-dependent.

The idiosyncratic patterns that were predictive of mind-wandering could be further appreciated via graph theoretical analyses and visualized maps of high-degree nodes (i.e., regions that participated in many predictive edges) (**Fig. 5d**). For example, these maps showed how in S3, high-degree nodes were particularly widely distributed, which explains why single-network retention CPMs consistently yielded significant performance. The top positive and negative nodes with ‘hub-like’ properties were different in each subject, as determined both based on node degree (**Table S3a**) and betweenness-centrality (**Table S3b**), though DMN-associated nodes were common across subjects. At the global level, S3 compared to the other two subjects showed higher integration across networks for both positive and negative CPM patterns, as determined both based on shorter characteristic path length (**Table S3c**) and its inverse, increased global efficiency (**Table S3d**).

### Cross-subject and population-derived models exhibit inconsistent performance

Our finding of individual differences in predictive network features hint that CPMs generated within individuals (or groups) may not always generalize to other individuals. To further examine this, we performed cross-subject CPM: for each pair of subjects, one subject was treated as training data and the other was treated as out-of-sample testing data. This analysis yielded inconsistent performance depending on the subject pair. Half of the train-test pairings resulted in significant cross-subject prediction of trial-wise mind-wandering (all *p* < 0.05, Spearman’s rank correlation) (**Fig. 6a**). Cross-subject CPMs between S2 and S3 resulted in bidirectional significant prediction, likely driven by overlapping predictive networks features between these subjects such as a negative association between mind-wandering and DMN-frontoparietal control network (FPCN) functional connectivity (**Fig. 5a**). Notably, these two subjects did not consistently report mind-wandering subcategories that were similar to one another, but they did demonstrate similarities in terms of emotional content and perspective-taking (**Fig. S1b**).

We next tested the generalizability of previously published, population-derived CPMs to the densely-sampled individuals investigated in the present study. We tested two population-derived CPMs of psychological constructs related to mind-wandering that have previously shown successful prediction of within-subject, trial-wise variability in stimulus-independent, task-unrelated thought (SITUT-CPM)^16^ and sustained attention (SA-CPM)^37^ (**Fig. 6b**). Applying the SITUT-CPM failed to yield significant out-of-sample predictions of mind-wandering in any of the densely-sampled subjects. Applying the SA-CPM revealed significant prediction of mind-wandering in only one of three subjects (*p* < 0.05, Spearman’s correlation). Taken together, cross-subject and population-level functional connectivity models resulted in inconsistent prediction of mind-wandering within densely-sampled subjects, highlighting the need for personalized models.

## Discussion

We combined dense-sampling fMRI, experience-sampling, precision functional mapping, and predictive modeling to investigate individual differences in the neural basis of self-reported mind-wandering (defined as off-task thought). We found reliable associations between DMN activation and mind-wandering when estimating networks within individuals and allowing for variable timing of spontaneous BOLD activation relative to subjective reports. Beyond DMN, large-scale networks activated and deactivated during mind-wandering were often distinct across individuals. Idiographic CPM revealed idiosyncratic network coupling patterns that consistently predicted mind-wandering within individuals, while cross-subject and population-derived models performed inconsistently when applied to densely-sampled individuals. Collectively, our findings detail previously unrecognized, substantial inter-individual variations in neural correlates and predictors of mind-wandering. We discuss the implications of our findings for conceptualizing mind-wandering and the brain, interpreting resting-state brain activity, and advancing clinical applications.

### Neural basis of mind-wandering: from populations to individuals

While it is recognized that mind-wandering is supported by multiple brain networks and neuromodulatory systems, the role of the DMN has received the most attention to date.^5,6,49^ Some of the first-described functional properties of the DMN were that this network deactivates during externally-oriented task performance^50,51^ and activates during mental experiences that are common during mind-wandering such as thinking about the past or future with self-related or social content.^52^ Neuroimaging studies,^53,54^ including those with online experience-sampling,^10–14^ subsequently offered evidence of a relationship between DMN activation and self-reported thoughts that are stimulus-independent and task-unrelated. Human intracranial recordings have further confirmed that neuronal population activity within DMN regions increases during attentional lapses.^55^ However, some fMRI studies have failed to show relationships between DMN activation and mind-wandering in group-level analyes,^15^ or in certain (non-densely-sampled) individuals,^13^ or have alternatively shown relationships with more specific features of thought such as subjective level of detail.^17^

We showed that when boosting within-subject statistical power, as well as accounting for personalized functional anatomy and BOLD temporal dynamics, DMN activation was reliably associated with mind-wandering (defined as off-task thought) during a simple visual fixation task. In two of three subjects (S1 and S3), DNa exhibited stronger associations than DNb. The functional differences between these two subnetworks remains an active research topic. One dense-sampling fMRI study showed that while DNa was recruited during remembering and imagining the future, DNb was alternatively recruited during social thoughts about others’ experiences (theory of mind).^35^ The subcategories of mind-wandering that our participants reported suggest that all individuals engaged in recalling, planning, and imagining (**Fig. S1a**), which may explain consistent DNa recruitment. While thought probes in this study did not inquire directly about theory of mind, we speculate that the more ‘equal’ recruitment of DNa and DNb found in one participant (S2) could suggest a tendency to engage in socially-oriented thoughts. Further, targeted investigations may address the distinct roles of DNa and DNb in spontaneous thought.

Our within-subject analyses revealed variability in the time courses of DNa and DNb activation relative to subjective reports, an effect that has not been appreciated in prior studies due to limited within-individual sampling. Interestingly, mind-wandering-related activation in these networks could sometimes be detected in BOLD activity up to ∼20 seconds prior to thought probe onsets (i.e., ∼25 seconds when accounting for hemodynamic delay). Relatedly, prior studies (at the group-level) have revealed trends of increased DMN activation^56^ and EEG alpha power^57^ up to ∼20-30 seconds prior to task performance errors. One potential interpretation of prolonged pre-probe DMN activation is that random-interval thought probes often capture mind-wandering when a person is already deeply immersed in self-generated experiences. In future studies, mind-wandering episodes may be captured more efficiently via real-time fMRI detection of DMN activation to potentially reduce the amount of within-subject sampling needed to identify brain-experience relationships.

Beyond DMN, we found that functional networks and regions most activated during mind-wandering were variable across subjects. One subject (S1) showed an activity pattern that largely aligns with expectations from population-level neuroimaging: the dorsal attention and salience networks, typically associated with externally-oriented attention,^58,59^ were most activated during task-focus, while DMN subnetworks were most activated during mind-wandering. This subject also showed mind-wandering-related hippocampal activation, potentially hinting at support for the hypothesis that hippocampal sharp-wave ripple events trigger DMN activation to initiate spontaneous thought.^60^ Another subject (S3) showed a whole-brain pattern that was not necessarily anticipated from population-level neuroimaging. The pattern included prominent activations of the language network and thalamic deactivations. As this subject reported a relatively high proportion of thoughts with auditory-language content (**Fig. S1a**), we speculate that thalamic deactivation reflected dampening of sensory input (i.e., perceptual decoupling^61^) to maintain attention toward self-generated inner speech. The third subject (S2) had weaker effect sizes than the others in terms of relationships between mind-wandering and activation of both the DMN and other networks. These weaker effects may be due to multiple factors, such as decreased statistical power (fewer analyzable trials available in this subject), ‘noisier’ introspective access to thought processes, or increased within-subject variability of thought subtypes.

We explored the broader role of intra- and inter-network interactions through network-based predictive modeling within individuals. This analysis revealed contributions of various intrinsic networks that were not necessarily captured in the BOLD activation analyses. This divergence is consistent with prior work demonstrating complementary, rather than overlapping, contributions of regional activation and functional connectivity to prediction of mind-wandering.^14,15^ Population-based predictive modeling analyses of mind-wandering have emphasized the importance of features that involve DMN within-network and between-network interactions,^14–16^ though broader interactions beyond DMN contributed substantially to the recently reported SITUT-CPM.^16^ Our densely-sampled subjects each showed idiosyncratic predictive networks that did not fully align with population-derived models. For example, while DMN-FPCN connectivity was positively related to mind-wandering in the SITUT-CPM,^16^ two subjects here showed a largely opposite pattern for DMN-FPCN edges. Though within-DMN features were sufficient for prediction within each subject, computational removal of the DMN had a minimal impact on prediction performance, and there were also other networks that were alone also sufficient for prediction performance. For example, the sensorimotor network, which has been previously linked to individual differences in mind-wandering content,^62^ was sufficient for prediction within all subjects.

One implication of our findings is there could be multiple neural ‘paths’ to mind-wandering (i.e., degeneracy). This possibility is in part supported by recent studies which demonstrated that EEG predictors of mind-wandering are distinct across individuals.^63,64^ Moreover, a recent application of fMRI multivariate pattern analysis revealed that person-specific, compared to population-derived, neural models could better decode affective valence when people reflected on concepts in spontaneous thought that were high in self-relevance.^29^ As degenerate mechanisms have been described at multiple levels of the nervous system,^65,66^ there is a need to more deeply investigate their potential importance to brain-experience relationships.

### Implications for interpreting spontaneous brain activity

Our findings build on research linking spontaneous brain activity to ongoing cognition.^67,68^ There has remained debate in the field over whether ‘thinking’, as contrasted with intrinsic physiological processes such as off-line plasticity and homeostasis, contributes meaningfully to functional network patterns that are observed in BOLD spontaneous activity (e.g. in “resting-state” fMRI).^42,69–72^ On the one hand, resting-state functional connectivity patterns are highly stable within individuals across multiple time scales.^70,73^ On the other hand, functional connectivity fluctuates on the order of seconds,^74,75^, and rapid fluctuations encode spontaneous behavior.^76^ Yet in typical resting-state fMRI studies, measures of ongoing cognition or behavior—especially of an individual’s unconstrained experiences—are rarely obtained.^77^

In a few prior works that involved limited individual-level sampling (e.g. single session fMRI), online experience-sampling during “rest” has revealed group-derived neural correlates of ongoing thought.^18,24^ Our findings advance upon those works, highlighting clearly, at the individual level, that dynamics of spontaneous BOLD activation and functional connectivity are relevant to reportable mental experiences. The effect sizes that we achieved with idiographic predictive modeling were often substantially higher than those typically reported for population-derived functional connectivity models of self-report outcomes,^16,78^ while we found that cross-subject and population-derived predictive models often failed when applied to individuals. These findings indicate that fluctuations in resting-state BOLD functional connectivity may more closely relate to ongoing experience than has been appreciated, yet such brain-experience relationships may not be detected when relying only on population-derived inferences.

Our findings may even suggest that functional connectivity ‘fingerprints’ (i.e., individual-specific, state-independent patterns^73^) are driven by tendencies to engage in certain forms of thought, which could be conceptualized as ‘thought fingerprints.’ Support for this idea comes from a recent study demonstrating that when spontaneous thoughts were abolished due to deep general anaesthesia, functional connectivity patterns became less inter-individually distinguishable.^79^ People often report certain patterns of thought that are relatively stable^80^ (despite influence of context^81^), which may imply that during resting states, thought trajectories tend to ‘default’ toward recurring patterns that an individual is predisposed to experiencing. These considerations promote the idea that resting-state fMRI interpretability may be enhanced through incorporating annotations of experiences.^42,72,77^

### Clinical implications of personalized neuroimaging

The limited group-to-individual generalizability illustrated here has implications for clinical efforts that aim to identify neuroimaging-based biomarkers or treatment targets. The way that ongoing thoughts tend to unfold for a given individual carries important consequences for mental health.^5^ Recently developed, population-derived neuroimaging predictive models of ongoing thought have been applied to patients diagnosed with psychiatric illness. The SITUT-CPM predicted excessive mind-wandering in adults with attention-deficit hyperactivity disorder.^16^ A resting-state fMRI model of trait rumination predicted depression scores in adults with major depressive disorder.^82^ However, such model predictions do not succeed in all individuals, and population-derived predictive models are most likely to fail in individuals who defy sample stereotypes (e.g. due to sociodemographic and/or clinical factors).^83^ As such, population-derived approaches are not likely to yield ‘one-size-fits-all’ biomarkers.^84^

An idiographic approach, as taken here, may offer an alternative way to identify state-related biomarkers of thought processes in a personalized manner. The potential clinical impact of this approach was illustrated in a recent *n*-of-1 study where intracranial electrophysiology was combined with experience-sampling in a patient with treatment resistant depression. A neural model of within-individual fluctuations in subjective mood was used to guide closed-loop deep brain stimulation and was predictive of the effects of stimulation on mood improvement.^85^ Idiographic neuroimaging models of spontaneous experience, as developed here, may similarly be used to guide personalized treatments in a non-invasive manner.

## Methods

### Participants and general procedure

Three study participants (S1, S2 and S3; age range: 25-29, 1 male) were recruited from the Duke-National University of Singapore (Duke-NUS) Medical School community. Each participant completed six 2-hour scanning sessions on different days (each separated by roughly one week). All participants had normal or corrected-to-normal vision and reported no neurological, psychiatric, or sleep disorders. Participants provided informed consent for procedures approved by the National University of Singapore institutional review board.

The full procedures of this study were reported elsewhere.^86^ Participants each completed six sessions of neuroimaging, which included functional localizer and experience-sampling conditions. Data analyses reported here are independent from those applied to this dataset previously, as prior work focused on a subset of experience-sampling trials that involved specific categories of visual and auditory imagery.^86^

### Neuroimaging data acquisition

Neuroimaging scans were performed with a 3-Tesla Siemens Prisma scanner equipped with a 12-channel head coil at Duke-NUS Medical School in Singapore. Functional magnetic resonance imaging (fMRI) runs were performed with a gradient echo-planar imaging multiband sequence (repetition time: 1.06 s, echo time: 32 ms, flip angle: 61°, field of view: 1980 x1980 mm, 2x2 mm in-plane resolution, 2mm slice thickness) with collection of thirty-six slices aligned with the AC-PC plane. Approximately 10 runs were collected per session for each participant. In each session, a T1-weighted anatomical image was also acquired (repetition time: 2.3 s, inversion time: 900 ms, flip angle: 8°, field of view: 256 x 240 mm, 192 slices, 1x1x1mm voxels).

### Experience-sampling paradigm

Participants completed 46 fMRI runs of a fixation task with experience-sampling across five neuroimaging sessions (7-11 runs per session; ∼5.8 hours totals). Before scanning, participants went through task training and instructions. The main task of the experiment was an 8-min fixation task with intermittent experience-sampling. Participants were instructed to stare at a fixation cross without thinking of anything in particular. Any thought unrelated to the task was defined as mind-wandering. Thought probes would appear at random intervals between 45-90 seconds. The first thought probe item inquired about the degree to which attention was focused on the task on a Likert scale of 1-8. Ratings between 1-4 indicated that their mind was wandering while ratings between 5-8 indicated task-focus. There was also an option to provide a “9” rating to indicate being distracted but not mind-wandering (e.g. attending to external stimuli such as scanner acoustic sounds). These “distracted” trials were removed from analyses. For the purposes of intuitive interpretation, we flipped the rest of the mind-wanderings ratings prior to all analyses (i.e., subtracted by 9) such that greater mind-wandering was associated with a higher rating.

If participants reported being focused on the task at the first probe, they were immediately brought back to the fixation task. Conversely, if participants indicated mind-wandering for the first thought probe item, then a series of additional items followed to elaborate on thought content and form. Thought probe items 2-5 probed (2) attentional awareness on scale of 1-8; (3) vividness on scale of 1-8; (4) perspective: first-person, third-person, or not sure; (4) thought type: recalling, planning, imagining, reasoning/thinking, or distracted; and (5) content description: visual, auditory, emotional, smell, taste, and bodily sensation (check all that apply). If categorization included visual thoughts, participants would next specify whether the imagery included faces, body parts, animals, plants, natural scenes, artificial scenes, motion, non-living objects, or food. If categorization included auditory thoughts, participants would specify whether the thought included animal sounds, language, music, or other sounds. Lastly, if categorization included emotional thoughts, participants would specify whether they were happy, surprised, angry, disgusted, sad, or afraid. There was no time limit for answering questions.

In addition to the pre-programmed thought probes, participants had the option of prompting probes on their own if they caught themselves mind-wandering before a probe appeared; however, there were no self-caught mind-wandering prompts reported by any of the three subjects. Intervals between probes were informed by the participants’ prior run performance in an attempt to catch one instance of mind-wandering every minute. If participants reported less mind-wandering with the initial interval settings in the previous block, the probe interval length was increased in the next block. Conversely, if participants reported more mind-wandering than anticipated, interval length was decreased.

During the task, eye-tracking was used to detect drowsiness. Prolonged eye closure twice in a session was interpreted as excessive drowsiness and the run was ended. The participant was then given a break before beginning the next run. Overall, 350 thought probes per subject appeared. Participants navigated probe response options using a button box in one hand, and they confirmed their selection(s) using another button box in their other hand.

### Functional localizer paradigms

Across 1 or 2 of the 6 sessions, participants each completed 10 total fMRI runs of functional localizer tasks (51.4 minutes total). These tasks were run for the original purpose of identifying regions of interest involved in visual and auditory processing.^86^ In the present study, the functional localizer data were used for the purpose of mapping personalized functional networks.

Retinotopic mapping scans were collected in three runs. In each run, stimuli of flashing checkerboard wedges were presented in different orientations across three conditions. These conditions were pseudo-randomized across six 20-s blocks with 20-s fixation preceding and succeeding each block. Across all fixation conditions, subjects were instructed to use buttons to indicate the color of the central cross as it switched between red and green. In the horizontal condition, two wedges appeared in-line horizontally beneath the fixation cross. In the vertical condition, another two wedges appeared in-line vertically beside the fixation cross. In the diagonal condition, four wedges appeared around the fixation cross along the diagonal axis. Each condition repeated twice per run.

Three additional runs of six 20-s blocks and 20-s fixation were performed for auditory mapping. In each block, successive tones from two of three frequency ranges, high (2370-5900 Hz), mid (880-2170 Hz), and low (340-870 Hz) were played in 10-s cycles. Within a cycle, the frequency of the tone was increased linearly for 5s before decreasing for another 5s. Subjects were instructed to press a button at the tone’s peak frequency.

Four runs of sixteen 20-s blocks were run to localize the visual regions involved in processing faces, scenes, and objects. Participants were presented with greyscale images of faces, scenes, common objects, and scrambled objects with four blocks dedicated to each category.

### Behavioral data analyses

Individual differences in experience-sampling ratings were summarized by visualizing relative proportions in mind-wandering content and frequency across all runs and sessions. Mind-wandering frequency was summarized by visualizing the relative proportion of mind-wandering trials to on-task trials. Mind-wandering trials were defined as trials where the response to probe question, “How focused were you on the task?” was greater than or equal to five (after reverse-coding). Mind-wandering content was summarized by visualizing the relative proportion of responses across subjects to the probe question, “Check all that well describes the contents of your mind-wandering” (**Fig. S1a**).

We examined whether mind-wandering ratings (response to first thought probe item) were associated with time-on-task (within runs and sessions) or head motion within each subject. Within-session time-on-task was defined as the ordered thought probe number within a session. Within-run time-on-task was defined as the onset time (in seconds) of a thought probe within a run. Head motion was defined as mean frame-wise displacement, as computed with the FMRIB software library (FSL) linear image registration tool,^87^ across the 28 repetition times (TRs) prior to each thought probe. We computed Spearman’s rank correlation coefficients between mind-wandering ratings and each of these metrics within subjects (significance set at *P* < 0.05, two-tailed).

To assess the degree to which subjects were similar to one another in their responses to thought probes, we performed a distance calculation on responses to categorical probe questions. The amount of times that a subject responded to each option for a given probe question was counted across all mind-wandering events. This yielded a vector of counts for each subject and for each probe question, where each element in the vector was a count of responses from a subject to a particular option for a probe question (e.g., [142, 79, 8] is the count vector of responses from subject 1 to probe question “Perspective” tallying the counts to options “First person”, “Third person”, and “Not sure”, respectively). The distance between two subjects for a probe question was compared by taking the Euclidean distance (i.e., square root of the sum of squared differences) between the respective count vectors. Lower distances suggest that two subjects responded more similarly to a given probe question throughout the experiment.

### fMRI preprocessing

Preprocessing of all 46 experience-sampling fMRI runs within each subject was performed using a combination of tools from FSL v6.0.3.,^88^ the analysis of functional neuroimages (AFNI v21.3.06) package,^89^ and independent components analysis for automatic removal of motion artifacts (ICA-AROMA). Procedures were similar to those previously described,^13^ though an exception was that we took extra steps to optimize within-subject alignment of all fMRI runs across sessions.^34^ We created a common BOLD template by computing a mean volume from all runs and sessions of a single participant. We then aligned each run to the common BOLD template using the MCFLIRT tool in FSL, thereby performing motion correction and moving all runs into the same common space. Using each subject’s T1 structural scan from their first MRI session, we performed segmentation with FSL’s FAST tool to create white matter (WM), cerebrospinal fluid (CSF), and grey matter (GM) templates. Using FSL’s FEAT to process fMRI data, we performed brain extraction (BET), spatial smoothing (5-mm full-width at half-maximum kernel), linear registration (FLIRT) between BOLD and T1 spaces, and nonlinear registration (FNIRT) between BOLD and standard MNI152 space. Motion-artifacts were then addressed with ICA-AROMA, which has been shown to improve sensitivity and specificity for both resting-state and task-based fMRI analyses.^90^ This included running ICA with automatic dimensionality estimation using FSL’s MELODIC tool and regression of automatically-detected motion-relevant components of fMRI data. Next, we set a threshold of 198 cm^3^ and 20 cm^3^ for WM and CSF probabilistic maps, deleted below-threshold volumes, and regressed out the mean WM and CSF time series from fMRI data. We applied a high-pass temporal filter (0.01 Hz cutoff) for univariate activation analyses or a bandpass filter (0.01 – 0.1 Hz) for functional connectivity analyses described below. The preprocessed data were transformed to MNI152 space using linear registration, and percent signal change (%SC) of voxel intensity values was computed by dividing the difference of values by the mean value in each run; the quotient was then multiplied by 100 for the final percent value. The functional localizer data were preprocessed with a different preprocessing pipeline to enhance consistency with pipelines that were used to validate the MS-HBM approach for personalized functional network mapping.^38^ We used publicly available fMRI preprocessing code shared by the Computational Brain Imaging Group (https://github.com/ThomasYeoLab/CBIG/tree/master/stable_projects/preprocessing/CBIG_fMRI_Preproc2016). Structural data (T1 scan from first session) were reconstructed into surface mesh representations using the recon-all command implemented in Freesurfer v6.0.0. Motion correction was performed using rigid body translation and rotation in FSL. Structural and functional images were aligned using boundary-based registration.^91^ We performed linear detrending and regressed out 9 regressors plus their temporal derivatives, including global signal, averaged WM signal, averaged CSF signal, six motion correction parameters. A bandpass filter of 0.01-0.1 Hz was applied. Preprocessed fMRI data were then projected to the Freesurfer fsaverage6 surface mesh, and spatial smoothing was performed on the surface with a 6mm full-width half-maximum kernel.

### Mapping personalized functional networks

We applied the MS-HBM approach^38^ to each subject’s functional localizer fMRI data to generate personalized functional connectivity networks, using publicly available code (https://github.com/ThomasYeoLab/CBIG/tree/master/stable_projects/brain_parcellation/Kong2019_MSHBM). While our functional localizer data involved multiple conditions, we deemed the combined data to be suitable for personalized network mapping because functional networks, as estimated over tens of minutes or more, show high stability within individuals regardless of task conditions.^70,92^ The MS-HBM method utilizes prior distributions of each cortical vertices’ functional connectivity profiles from various hierarchies (e.g. group-level, inter-subject level, intra-subject inter-session level) to estimate individual-specific brain networks unique to each subject.^38^

Model priors were computed based on initial binarized functional connectivity profiles calculated for multiple equally spaced cortical vertices, or designated points across the surface of the cortex. These functional connectivity profiles were defined as the top 10% of all correlations between the preprocessed BOLD signal at each cortical vertex and those of a predefined set of 1175 regions spanning the cerebral cortex. To estimate the first prior hierarchy (a group-level network profile), the average binarized connectivity profiles of all cortical vertices of all subjects was computed. This was then used to model the next prior (inter-subject network profile) for each subject, which considers the functional connectivity variability between subjects. The following prior (intra-subject inter-session network profile) compares the average connectivity profiles of all vertices from one session of a subject to those of that subject’s other sessions. Finally, the connectivity variance between different regions of the same network as observed in a single session of a subject (intra-session inter-region profile) was modeled.

These priors then informed the model to stabilize estimations of individually parcellated whole-brain networks, such that if the connectivity profile of a given vertex is most similar to the connectivity profile of a certain brain network at a specific prior hierarchy, the vertex would then be assigned to that network. For each subject, each cortical vertex was assigned to one of *k*=17 networks (where *k* is number of clusters), consistent with the 17-network solution of the population-level Yeo atlas.^40^ This approach of individually parcellating brain networks allowed us to analyze each subject’s brain activity within their own personalized cortical map. To inspect the consistency of networks obtained with MS-HBM in our dataset, we repeated the procedures with the algorithm constrained to *k* values of 13-16.

### Region-of-interest analyses of default mode network

We visually inspected the 17 networks obtained with MS-HBM within each subject and identified the personalized networks that were most consistent with the previously described DMN subnetworks known as “DNa” and “DNb”.^34,39^ We projected the network masks from fsaverage6 surface space to the MNI152 volumetric space of the preprocessed experience-sampling fMRI data. We then extracted the median BOLD %SC time series, separately from DNa and DNb voxels. For each thought probe across all runs and sessions, we computed BOLD %SC within each network averaged across 9 TRs prior to thought probes (i.e., corresponding to 9.54 seconds). We computed Spearman’s rank correlation coefficient between trial-wise mind-wandering ratings and the ∼10-second preprobe BOLD %SC values for the DNa and DNb (significance set at *P* < 0.05, two-tailed). We then repeated these analyses using alternative, standard-space DMN subnetworks instead of the personalized DNa and DNb subnetworks. For this comparison analysis, we extracted BOLD %SC from three DMN subnetworks in the Yeo17 population-level atlas,^40^ which have been described as “Core DMN,” “dorsomedial prefrontal cortex subsystem,” and “medial temporal lobe subsystem.”^41^

Beyond our analyses that focused on averaged BOLD %SC within ∼10-second windows prior to thought probes, we performed a complementary analysis of TR-by-TR temporal dynamics within DNa and DNb. For each subject, we split trials into “pure” high mind-wandering (rating between 6-8) and non-mind-wandering (rating between 1-3) while discarding trials that had intermediate ratings. At each TR between -20.14 and 0 seconds prior to thought probes (i.e., 20 TRs), we computed the average of median BOLD %SC within DNa and DNb across trials for each trial type. At each of these 20 TRs (and for each subnetwork), we performed Wilcoxon rank-sum tests to compare BOLD %SC between high and low mind-wandering trials. Significance was set at *P* < 0.05 (two-tailed), false-discovery rate corrected for multiple comparisons across 20 TRs. For visualization purposes, we further plotted average time courses of high and low mind-wandering trials prior to and after the ∼20 second window of interest.

### Whole-brain general linear model analyses

We performed within-subject, voxel-wise general linear model (GLM) analyses in FSL to search the whole brain for regions associated with mind-wandering ratings. For each experience-sampling fMRI run, we ran a first-level GLM on preprocessed data using FMRIB’s improved linear model prewhitening. This model included a parametric regressor for mind-wandering rating at the 10-second period prior to thought probes (convolved with a gamma hemodynamic response function). Contrasts were performed to identify voxel clusters associated positively or negatively with mind-wandering rating. Due to lack of variation in mind-wandering ratings within a small number runs (S1: 0 runs; S2: 1 run; S3: 1 run), those runs were excluded from GLM analyses. The first-level GLM statistical maps were submitted to higher-level, within-subject GLMs that combined all runs and sessions and were performed with FMRIB’s local analysis of mixed effects (FLAME) 1+2 (cluster-based thresholding at *Z* > 3.1 and family-wise error corrected *P* < 0.05). For visualization purposes, GLM results were projected to the fsaverage6 cortical surface and displayed with Connectome Workbench software.^93^ To provide further interpretation of whole-brain GLM results in terms of personalized cortical networks, we extracted the median *Z* statistic score within each network identified by MS-HBM.

### Within-subject connectome-based predictive modeling

We performed idiographic (within-subject) CPM,^94^ using methods consistent with those reported previously.^16^ We used both leave-one-trial-out and five-fold cross-validation to verify the stability of results. To generate features for predictive modeling, we extracted BOLD preprocessed time series from 300 regions in the Schaefer atlas from each fMRI experience-sampling run. We then correlated the time series between all region-pairs and Fisher *z*-transformed the Pearson correlation coefficients to generate a functional connectivity matrix for each trial within the 30-s window (28-frames) preceding the appearance of each thought probe.^16^ For each cross-validation fold, we identified edges (pairs of regions) with a correlation, positive or negative, to mind-wandering ratings that surpassed an uncorrected threshold of *P* < 0.01 (two-tailed). This step generated a “positive mask” and “negative mask,” each comprised of edges identified as features associated with mind-wandering. For each trial, we computed the dot product between the functional connectivity matrix and each mask. We then summed all positive and negative mask edge values. We calculated single network strength value based on the difference between summed negative mask and summed positive mask edge values. Using all trials within a given fold, we fit a linear model with network strength as the independent variable and mind-wandering as the dependent variable. This model was then applied to predict mind-wandering for the held-out trial(s), and predicted ratings were correlated with observed mind-wandering ratings. For five-fold cross-validation, the procedures described above were repeated 120 times with randomized cross-validation fold schemes, and the mean correlation between predictive versus observed mind-wandering was computed across iterations. To determine statistical significance of CPM, we performed 1000 permutation tests to generate null correlation values for predicted versus observed mind-wandering. Significance was set at *P_perm_* < 0.05.

In control analyses, we repeated the CPM procedures but during model training, we performed partial correlations between predictive versus observed mind-wandering, controlling for the following variables: (1) time-on-task within run; (2) trial number within session; and (3) head motion (i.e., mean framewise displacement within 30-second window prior to thought probe). For null distribution computation (based on 1000 permutations), mind-wandering ratings and control variable values were jointly shuffled.

Recent work has suggested that alternative predictive modeling methods may outperform CPM in producing reliable functional connectivity patterns.^48,95^ To ensure that our results were not limited by the CPM approach, we performed brain basis set modeling,^96^ which reduces input features (i.e., edge) using principal components analysis before conducting linear regression of the outcome scores. We chose to reduce the data to 75 components as per previous studies.^48,95^ Significance was assessed with permutation tests as done in the CPM analyses.

### Analyses of CPM feature importance

To interpret each individualized mind-wandering CPM, we generated positive and negative CPM masks within each subject using a single fold that included all trials (i.e., we correlated mind-wandering rating versus functional connectivity for each edge and set an uncorrected threshold of *P* <0.01, two-tailed, for positive and negative masks). We mapped the positive and negative CPM mask edges on to the Schaefer300 atlas, with each node assigned to one of seven standard Yeo-Krienen networks.^40^ We quantified the number of edges belonging to each intra- or inter-network pair (28 total network pairs).

Next, we performed statistical tests of feature importance using CPM with computational network lesions. We performed these analyses with two variations: (1) Single-network lesion: all edges assigned to one Yeo-Krienen network were removed during CPM training and testing; and (2) Single-network retention: all edges that were not assigned to a specific Yeo-Krienen network were removed during CPM training and testing. Computationally lesioning all but one of the seven Yeo-Krienen networks, we generated individualized subject CPMs based on each of the isolated seven networks to determine which, if any, network, could predict mind-wandering independently. Statistical significance was assessed in the same manner as in the main CPM analyses with 1000 permutation tests and significance set at *P_perm_* < 0.05.

To further illustrate individual variability in features contributing to CPMs, we plotted node degree using the BioImage Suite Connectivity Visualization Tool (https://bioimagesuiteweb.github.io/webapp/connviewer.html). These plots were based on the main within-subject CPM analyses performed using the Shen atlas of 268 regions^47^ rather than the Schaefer 300 atlas (CPM performance was very similar regardless of atlas choice). The node degree plots illustrate, for each node in the atlas, the total number of edges that the node participated in for positive and negative CPM masks (**Fig. 5d**).

We computed graph theory measures from the positive and negative networks obtained from CPM, using the Brain Connectivity Toolbox (https://sites.google.com/site/bctnet/).^97^ The networks obtained from CPM can be treated as binary undirected networks. We calculated global measures of integration including efficiency and characteristic path length. These measures assess the degree to which nodes communicate. We also calculated local measures of node centrality, or the importance of a node for efficient communication. Specifically, we calculated betweenness-centrality, the fraction of shortest paths that pass through a node, and degree, the number of connections involving the node.

### Cross-subject CPM

We tested model generalizability across subjects using cross-subject testing of the personalized models generated by idiographic CPM. To obtain within-subject CPMs that could be tested out-of-sample in other subjects, we generated a CPM linear model within each subject using a single fold that included all trials. We applied the linear model from each individualized CPM to generate predicted mind-wandering ratings for the remaining two subjects (i.e., if using the CPM trained on subject 1, the linear model was applied to subjects 2 and 3 respectively). In external data from test subjects, we computed network strength based on the dot products of trial-wise functional connectivity matrices and the positive and negative edge masks from the model training subject. Network strength was computed based on subtraction of negative from positive edge scores. We computed predicted mind-wandering scores based on linear parameters obtained in the model training subject. We then calculated the Spearman’s rank correlation coefficient for the predicted versus observed mind-wandering ratings (significance set at *P* < 0.05, two-tailed).

### Testing population-derived CPMs

We tested the ability of two previously published, population-derived CPMs (i.e., those that combined features derived from multiple individuals during model training) to predict mind-wandering within each of our densely-sampled subjects. These two CPMs included: (1) The stimulus-independent and task unrelated thought CPM (SITUT-CPM),^16^ publicly available at https://github.com/swglab/CPM_CONN; and (2) The sustained attention CPM (SA-CPM),^98^ publicly available at https://github.com/monicadrosenberg/Rosenberg_PNAS2020/tree/master. We tested the SITUT-CPM because this model was predictive (at a group-level) of within-subject mind-wandering away from a continuous performance task, both within healthy adults and adults diagnosed with attention-deficit hyperactivity disorder.^16^ The SA-CPM was shown to be predictive of sustained attention ability, both across^98^ and within^37^ individuals, and showed a (weak) negative correlation with SITUT-CPM expression.^16^

To test the population-derived CPMs, we computed trial-wise network strength based on the dot products of trial-wise functional connectivity matrices and the positive and negative edge masks from the respective CPM. Trial-wise network strength was computed based on subtraction of negative from positive edge scores. We then computed the Spearman’s rank correlation coefficient between CPM network strength and observed mind-wandering ratings (significance set at *P* < 0.05, two-tailed). To test the SITUT-CPM, features were derived from the Schaefer 300-region atlas (as used in the idiographic CPM analyses). The SA-CPM was created based on the Shen atlas of 268 regions,^47^ and so the same atlas was used to extract features when testing this model.

## Acknowledgments and Funding Sources

This work was supported by the National Institute of Mental Health of the National Institutes of Health under award number R21MH129630 (to AK) and T32NS047987 (to NA).

## Author contributions

A.K.: Conceptualization, methodology, software, formal analysis, investigation, visualization, writing-original draft, writing-review and editing. N.A.: Formal analysis, methodology, software, writing-reviewing and editing. T.B.: Formal analysis, visualization, writing-original draft, writing-reviewing and editing. D.B.: Formal analysis, visualization, writing-original draft, writing-reviewing and editing. L.S.T.: Formal analysis, writing-original draft, writing-reviewing and editing. Braga, R.M.: Formal analysis, methodology, writing-reviewing and editing. S.H.: Conceptualization, investigation, methodology, writing-reviewing and editing.

## Supplementary Information

**Figure S1.**
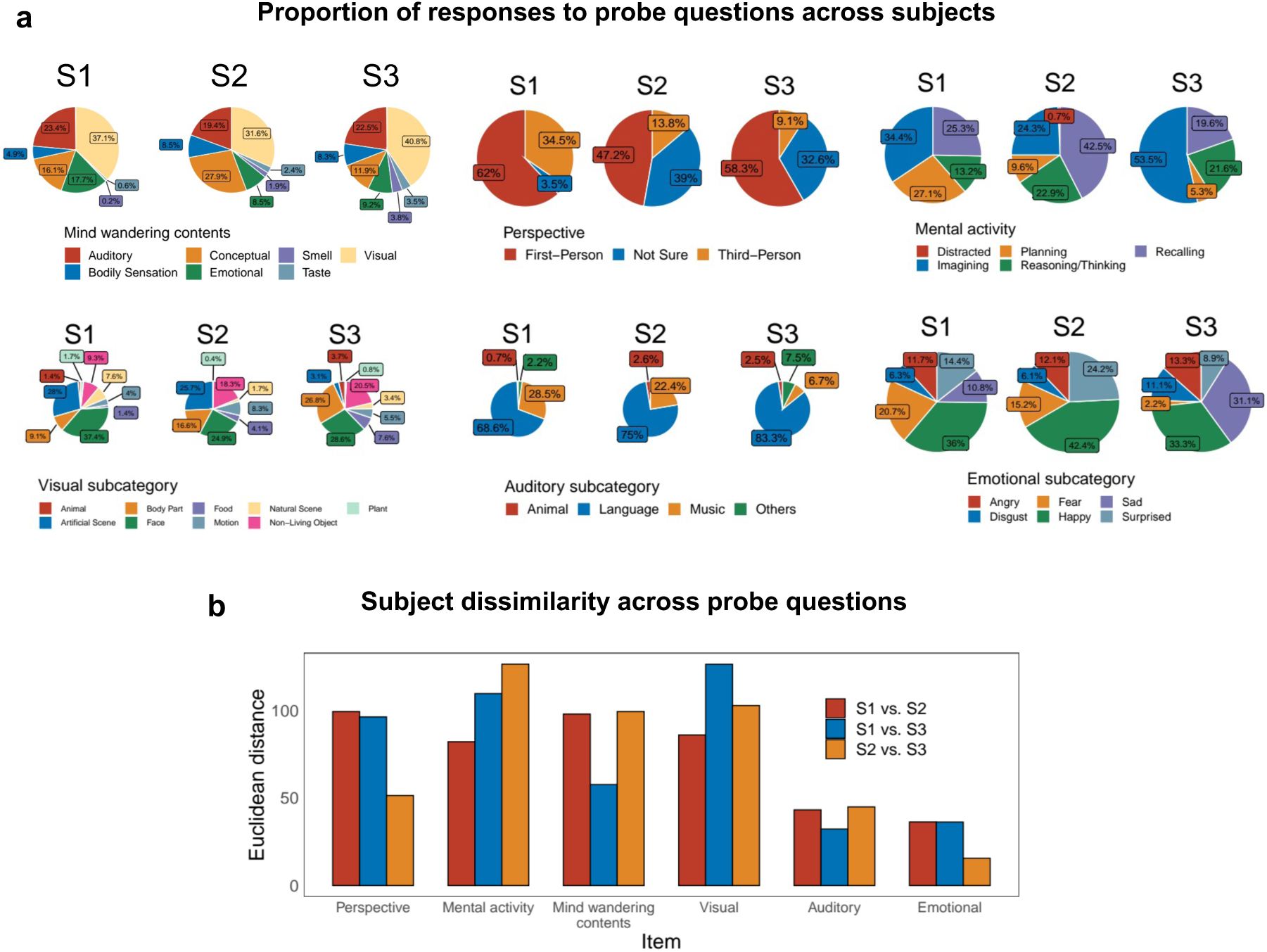
Individual differences in responses to mind wandering subcategory probes. **a**) For each subject, proportion of responses to each option for six probe questions concerning contents, perspective, type of mental activity, visual subcategory, auditory subcategory, and emotional content. **b**) Euclidean distance between response counts for each pair of subjects and each probe question. The distance between two subjects for a probe question was computed by first tallying the number of responses to each option for each subject, and then computing the Euclidean distance between the two tallies.

**Figure S2.**
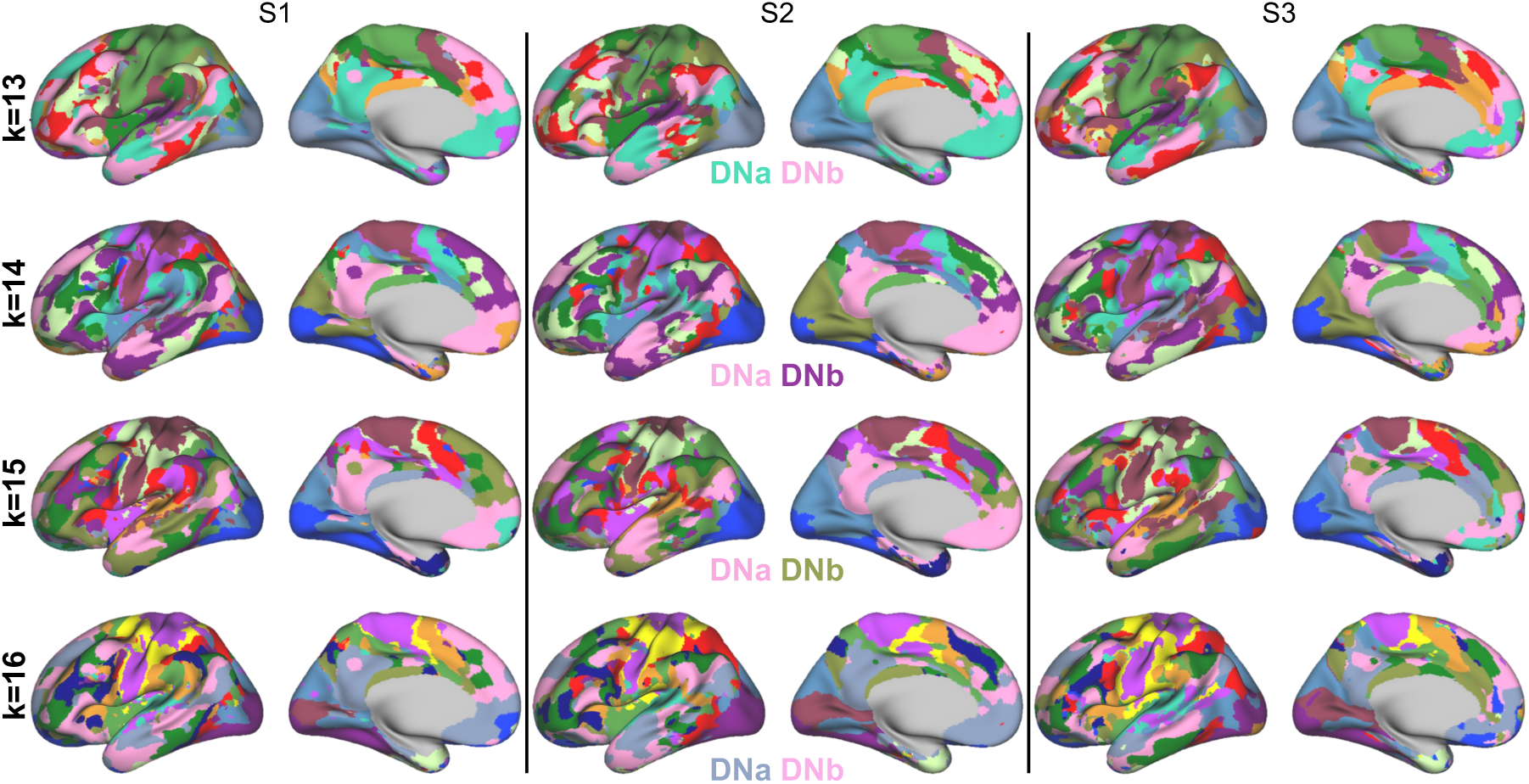
Personalized functional networks estimated with multi-session hierarchical Bayesian modeling (MS-HBM) with multiple clustering solutions. For each subject and clustering solutions set between 13 and 16 (see Fig. 3 for 17 cluster solution), cortical networks obtained with MS-HBM are shown. Consistency was found across solutions in identifying two subnetworks of the default mode network, default network A (DNa) and B (DNb).

**Figure S3.**
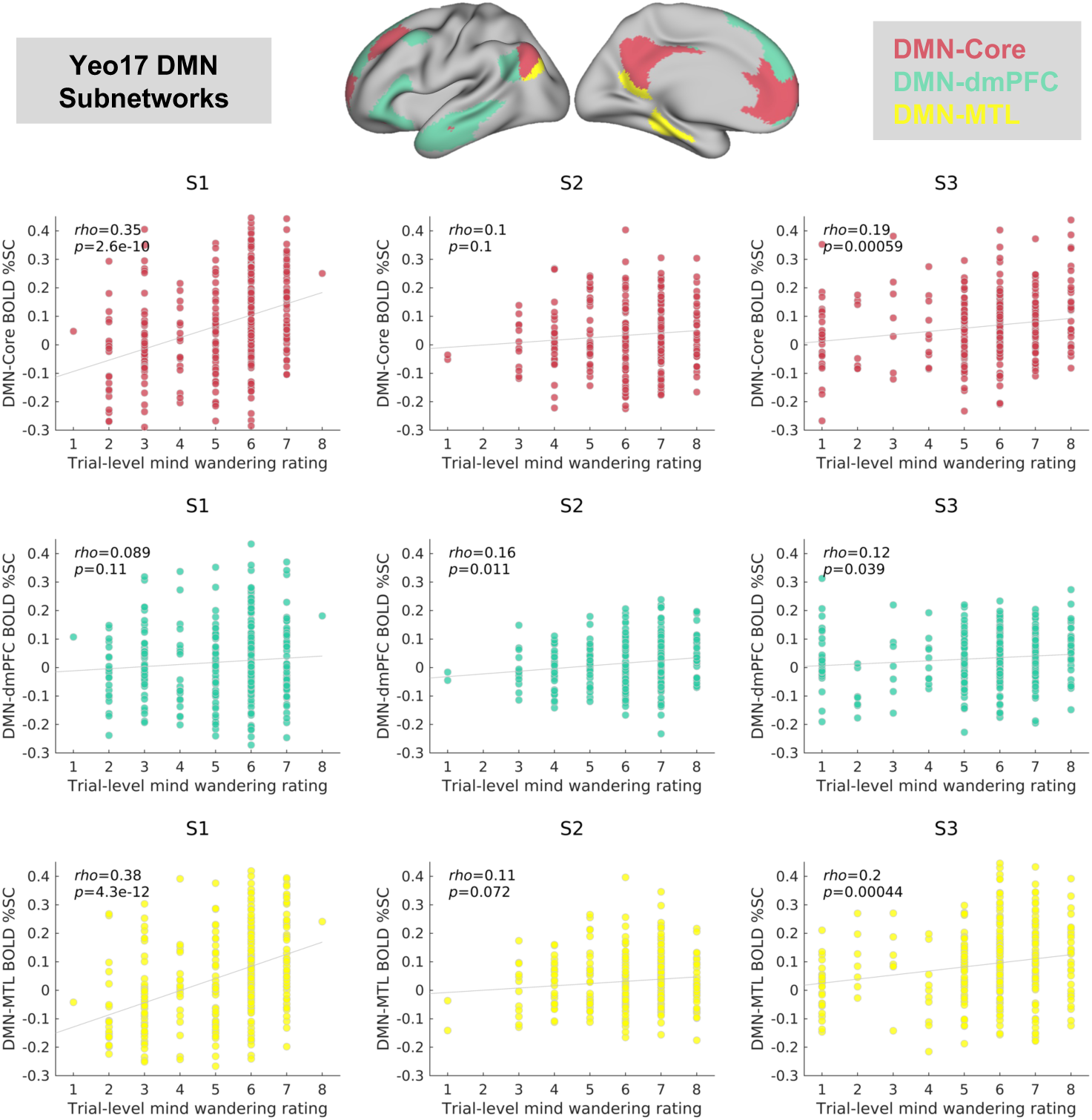
Correlations between mind wandering and activation within standard-space subnetworks of the default mode network (DMN). The top cortical surface plots illustrate the locations of standard-space subnetworks from the population-derived Yeo-Krienen 17-network atlas. These networks include the DMN-core, DMN-dorsomedial prefrontal cortex (dmPFC) and DMN-medial temporal lobe (MTL) subsystems. Scatter plots show correlations within each subject between trial-by-trial mind wandering and the median of blood oxygenation level dependent (BOLD) percent signal change (%SC) within 10-second pre-thought probe periods for a given DMN subnetwork. The top, middle and bottom rows of plots, respectively, show results for the DMN-Core, DMN-dmPFC, and DMN-MTL subnetworks.

**Table S1.**
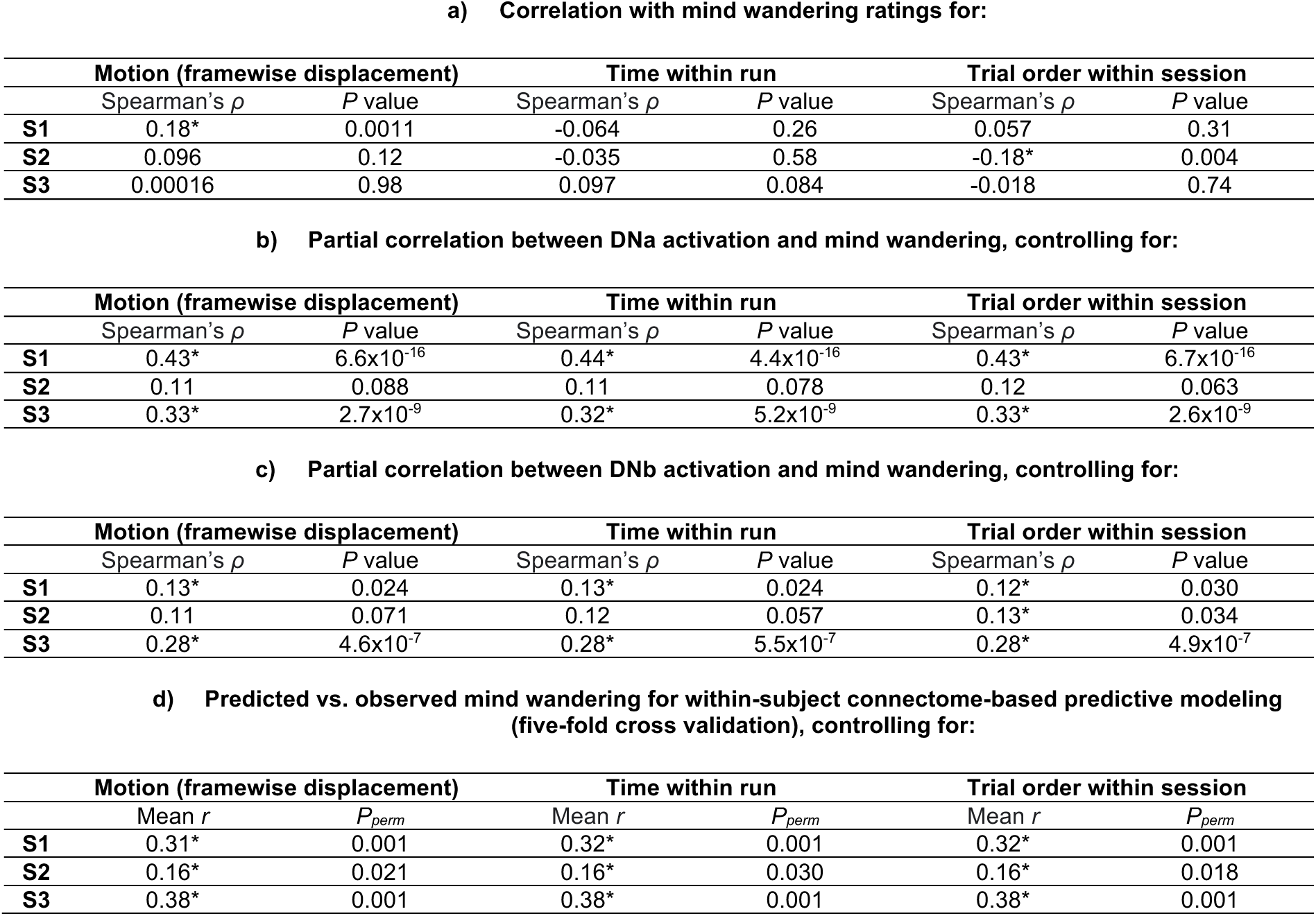
Control analyses accounting for head motion and time on task. **a)** Correlations between trial-wise mind wandering and head motion (i.e., mean framewise displacement in 30-second windows prior to thought probes), time within run (i.e., onset time of thought probe from start of each run), and trial order within session for all subjects. **b)** Partial correlations between trial-wise mind wandering and DNa activation (BOLD %SC in 10-second windows prior to thought probes) controlling for motion and time on task. **c)** Same as b) but for DNb instead of DNa. **d)** Results of within-subject connectome-based predictive modeling of mind wandering (five-fold cross validation), using partial correlations within each cross-validation fold to account for head motion and time-on-task. Mean *r* values indicate the average of predicted versus observed mind wandering correlations across 120 cross-validation iterations, and *P* values are based on 1000 permutations. \**P*<0.05. DNa = default network A; DNb = default network B.

**Table S2.** Alternative predictive modeling results. **a)** Results of within-subject connectome-based predictive modeling of mind wandering (five-fold cross validation), using the Schaefer 300-region and Shen 268-region atlases. **b)** Results of within-subject brain basis set (BBS) modeling of mind wandering (five-fold cross validation), using the Schaefer 300-region and Shen 268-region atlases. \**P*<0.05

|  | Schaefer300 |  | Shen268 |  |
| --- | --- | --- | --- | --- |
| | Mean $r$ | $P_{perm}$ | Mean $r$ | $P_{perm}$ |
| **S1** | 0.31* | 0.001 | 0.31* | 0.001 |
| **S2** | 0.16* | 0.025 | 0.20* | 0.008 |
| **S3** | 0.38* | 0.001 | 0.39* | 0.001 |

b) BBS
|  | Schaefer300 |  | Shen268 |  |
| --- | --- | --- | --- | --- |
| | Mean $r$ | $P_{perm}$ | Mean $r$ | $P_{perm}$ |
| **S1** | 0.38* | 0.001 | 0.27* | 0.001 |
| **S2** | 0.11 | 0.074 | 0.22* | 0.007 |
| **S3** | 0.29* | 0.001 | 0.34* | 0.001 |

**Table S3.** Graph theory metrics for features selected by connectome-based predictive models in each subject. **a)** Top region (and associated network labels from Schaefer 300-region atlas) with highest node degree in each subject, for positive and negative CPM networks respectively. **b)** Top region (and associated network label from Schaefer 300-region atlas) with highest betweenness centrality (BC) in each subject, for positive and negative CPM networks respectively. **c)** Characteristic path length (CPL) in positive and negative CPM networks. Smaller CPL indicates greater global integration. **d)** Global efficiency in positive and negative CPM networks. Greater global efficiency indicates greater integration. DMN = default mode network; SAL = salience network. SMN = sensorimotor network.

**a) Node degree**
|  | Positive Network |  |  | Negative Network |  |  |
| --- | --- | --- | --- | --- | --- | --- |
|  | Region Label | Network Label | Degree Value | Region label | Network Label | Degree Value |
| **S1** | Left temporal pole | Limbic | 36 | Right temporal-parietal junction | SAL | 71 |
| **S2** | Left precuneus/posterior cingulate cortex | DMN | 17 | Left somatomotor cortex | SMN | 45 |
| **S3** | Left pregenual anterior cingulate cortex | DMN | 121 | Left somatomotor cortex | SMN | 138 |

**b) Node betweenness centrality**
|  | Positive Network |  |  | Negative Network |  |  |
| --- | --- | --- | --- | --- | --- | --- |
|  | Region Label | Network Label | BC value | Region Label | Network Label | BC Value |
| **S1** | Left temporal pole | Limbic | 10974 | Right precuneus/posterior cingulate cortex | DMN | 6049 |
| **S2** | Left precuneus/posterior cingulate cortex | DMN | 2209 | Right orbitofrontal cortex | Limbic | 6745 |
| **S3** | Left pregenual anterior cingulate cortex | DMN | 8782 | Right frontal pole | DMN | 5298 |

**c) Characteristic path length**
|  | Positive Network | Negative Network |
| --- | --- | --- |
| **S1** | 3.97 | 2.87 |
| **S2** | 4.79 | 3.00 |
| **S3** | 2.47 | 2.34 |

**d) Global efficiency**
|  | Positive Network | Negative Network |
| --- | --- | --- |
| **S1** | 0.29 | 0.40 |
| **S2** | 0.28 | 0.37 |
| **S3** | 0.45 | 0.52 |

